# Convergent losses of *TLR5* suggest altered extracellular flagellin detection in four mammalian lineages

**DOI:** 10.1101/2020.02.23.962035

**Authors:** Virag Sharma, Felix Walther, Nikolai Hecker, Heiko Stuckas, Michael Hiller

**Author notes:** To whom correspondence should be addressed, Michael Hiller, Computational Biology and Evolutionary Genomics, Max Planck Institute of Molecular, Cell Biology and Genetics & Max Planck Institute for the Physics of Complex Systems, Dresden, Germany. current affiliations: CRTD-DFG Center for Regenerative Therapies Dresden, Carl Gustav Carus Faculty of Medicine, Technische Universität Dresden, Dresden; Paul Langerhans Institute Dresden (PLID) of the Helmholtz Center Munich at University Hospital Carl Gustav Carus and Faculty of Medicine, Technische Universität Dresden, Dresden; German Center for Diabetes Research (DZD), Munich, Neuherberg, Germany.

## Abstract

Toll-like receptors (TLRs) play an important role for the innate immune system by detecting pathogen-associated molecular patterns. *TLR5* encodes the major extracellular receptor for bacterial flagellin and frequently evolves under positive selection, consistent with coevolutionary arms races between the host and pathogens. Furthermore, *TLR5* is inactivated in several vertebrates and a *TLR5* stop codon polymorphism is widespread in human populations. Here, we analyzed the genomes of 120 mammals and discovered that *TLR5* is convergently lost in four independent lineages, comprising guinea pigs, Yangtze river dolphin, pinnipeds, and pangolins. Validated inactivating mutations, absence of protein-coding transcript expression, and relaxed selection on the *TLR5* remnants confirm these losses. PCR analysis further confirmed the loss of *TLR5* in the pinniped stem lineage. Finally, we show that *TLR11*, encoding a second extracellular flagellin receptor, is also absent in these four lineages. Independent losses of *TLR5* and *TLR11* suggests that a major pathway for detecting flagellated bacteria is not essential for different mammals and predicts an impaired capacity to sense extracellular flagellin

## Introduction

Toll-like receptor (TLR) proteins are an important component of the innate immune system (Vijay 2018). TLRs comprise a group of membrane-bound pattern recognition receptors and detect pathogen-associated molecular patterns such as bacterial lipopolysaccharides, peptidoglycans, double-stranded RNA or flagellin (Vijay 2018). Extracellular flagellin is recognized by TLR5 (toll-like receptor 5) (Hayashi, et al. 2001; Mizel, et al. 2003; Smith, et al. 2003). Upon binding its ligand, TLR5 stimulates the expression of proinflammatory, antibacterial and stress-related genes (Yu, et al. 2003; Vijay-Kumar, et al. 2008), and hence plays an important role for recognizing pathogenic flagellated bacteria such as Salmonella (Murthy, et al. 2004). The flagellin sensor function of TLR5 is likely conserved among vertebrates given that TLR5 of chicken, Anole lizard and rainbow trout also recognizes bacterial flagellin (Tsujita, et al. 2004; Iqbal, et al. 2005; Voogdt, et al. 2016). Importantly, *Tlr5* knockout mice do not show an immune response to injected flagellin (Feuillet, et al. 2006; Uematsu, et al. 2006; Vijay-Kumar, et al. 2007), suggesting that Tlr5 is the main extracellular flagellin sensor.

Coevolutionary arms races between pathogens and the host likely explain why pathogen-sensing immune genes frequently evolve under positive selection (Sackton, et al. 2007). Consistently, the *TLR5* gene experienced positive selection in many primate, mammal and bird lineages, and recurrent positive selection was often detected for sites located near the ligand binding site (Wlasiuk, et al. 2009; Alcaide, et al. 2011; Areal, et al. 2011; Smith, et al. 2012; Vinkler, et al. 2014; Velova, et al. 2018). Interestingly, despite its conserved role in detecting flagellin, previous studies revealed that *TLR5* is inactivated (lost) in several independent bird lineages (Alcaide, et al. 2011; Bainova, et al. 2014; Velova, et al. 2018), in tuatara and in clownfish (Liu, et al. 2019). Furthermore, a dominant negative *TLR5* polymorphism that introduces a premature stop codon is present in ~4% of human individuals and has reached a frequency of >20% in some populations (Hawn, et al. 2003; Barreiro, et al. 2009; Wlasiuk, et al. 2009). Therefore, the question arises whether inactivation of *TLR5* also occurred in non-human mammals. Here, by analyzing the genomes of 120 mammalian species, we show that *TLR5* is lost in four independent mammalian lineages, comprising guinea pigs, Yangtze river dolphin, pinnipeds, and pangolins.

## Results

We applied a previously developed approach (Sharma, et al. 2018a) to screen for gene-inactivating mutations in *TLR5* using a genome alignment of 120 placental mammals (Supplementary Table 1) (Hecker, et al. 2020). This screen revealed that the single coding exon of *TLR5* exhibits stop codon mutations, smaller frameshifting insertions or deletions, or larger deletions in the genome assemblies of 17 species belonging to 14 distinct lineages (gorilla, orangutan, guinea pig, rabbit, pig, alpaca, Yangtze river dolphin, bighorn sheep, Weddell seal, Hawaiian monk seal, walrus, Malayan and Chinese pangolins, little brown bat, elephant, armadillo). Since we have previously shown that apparent gene-inactivating mutations can sometimes be sequencing or assembly errors (Hecker, et al. 2017; Sharma, et al. 2018b; Hecker, et al. 2019a; Huelsmann, et al. 2019), we carefully validated the correctness of all such putative mutations and performed additional analyses to establish which of these 17 species truly lost *TLR5*.

First, we used available raw sequencing read data to validate stop codon and smaller frameshifting insertion/deletion mutations. This revealed that frameshifting insertions and deletions that are apparent in the orangutan, rabbit, alpaca, bighorn sheep, little brown bat, elephant and armadillo assemblies are in fact base errors in the assemblies as aligning raw sequencing reads do not validate these putative mutations (Supplementary Figure 1A-J). Similarly, sequencing reads show that 1 bp frameshifting deletions in the gorilla (gorGor5) and pig (susScr11) genomes that are assembled from PacBio long sequencing reads (Gordon, et al. 2016; Warr, et al. 2019) are base errors (Supplementary Figure 2). Furthermore, these two base errors are not present in previous assemblies of gorilla (gorGor3) and pig (susScr3), and therefore are likely due to uncorrected errors present in raw PacBio reads (Watson, et al. 2019). In contrast to these nine species, all frameshifts and stop codon mutations present in the domestic guinea pig, Weddell seal, and Pacific walrus were confirmed by several sequencing reads and we found no support for the ancestral, non-inactivating allele (Figure 1A, Supplementary Table 2).

**Figure 1:**
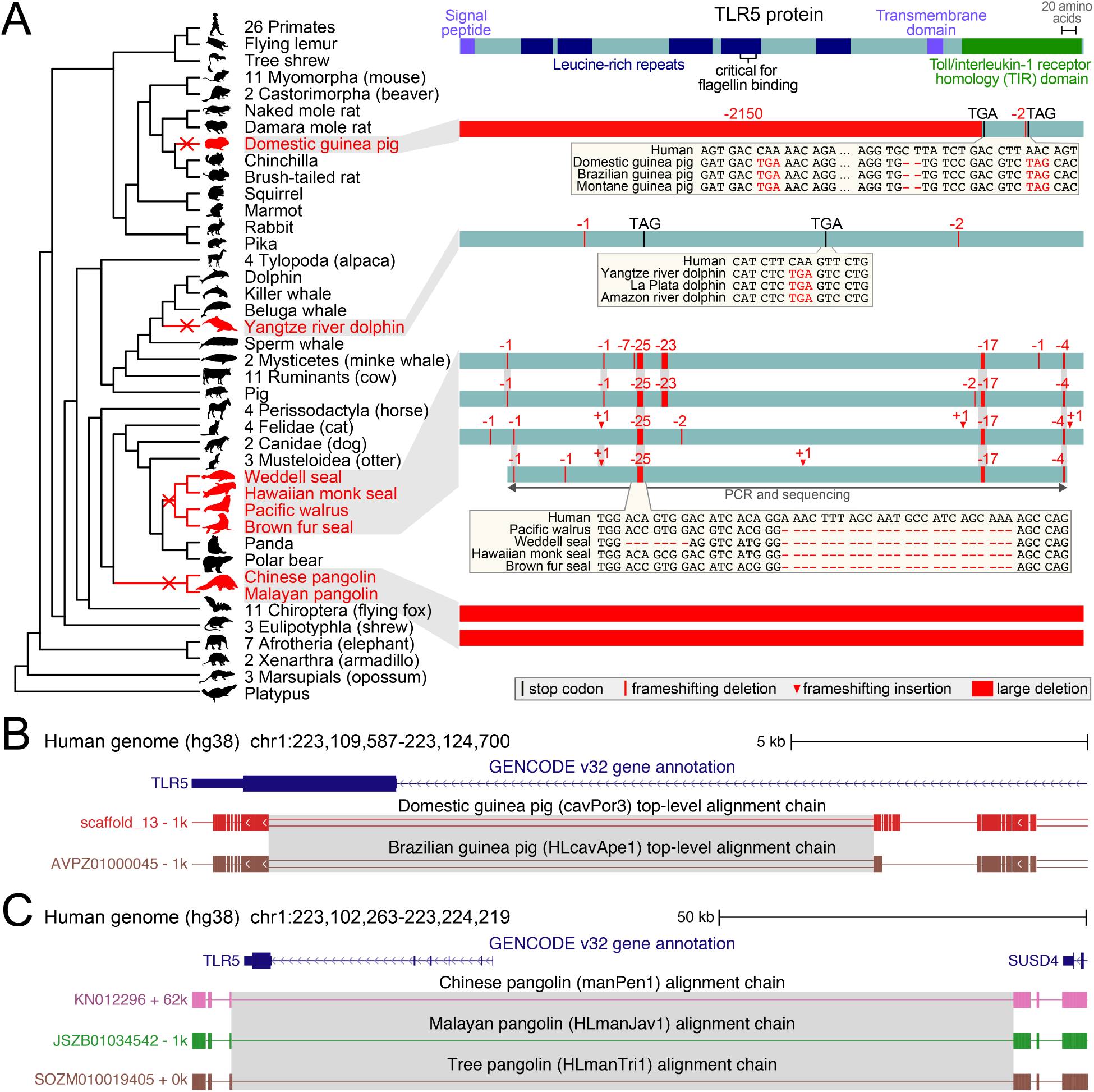
Loss of *TLR5* in four independent mammalian lineages. (A) Left: Phylogeny of mammals, showing species that lost *TLR5* in red font. Right: The human TLR5 protein is visualized at the top, superimposed with the signal peptide, transmembrane and Pfam protein domains, as annotated in Ensembl (Yates, et al.), and the region required for flagellin binding (Mizel, et al. 2003). Boxes below represent the *TLR5* coding exon of individual species, superimposed with gene-inactivating mutations. For the brown fur seal, we sequenced the majority of the *TLR5* exon. Insets show inactivating mutations (red font) that are shared among sister species. Mutations shared between at least two pinniped species are highlighted by grey background. (B) UCSC genome browser (Haeussler, et al. 2019) view of the human hg38 genome, showing the *TLR5* gene and top-level chain of co-linear local alignments (blocks represent aligning regions, double lines represent unaligning sequence, and single lines represent deletions) to two guinea pig species. Alignment chains show that the large deletion that removes the majority of the *TLR5* coding exon (grey background) has the same breakpoints and thus likely occurred in the ancestor of both guinea pigs. (C) UCSC genome browser view shows that the *TLR5* deletion has breakpoints shared between three pangolin species, suggesting that *TLR5* loss already occurred in their common ancestor.

Second, we analyzed the pairwise genome alignments between human and guinea pig, river dolphin, pinnipeds, and pangolins. This confirmed that the remnants of *TLR5* occur in a genomic locus with conserved gene synteny (Supplementary Figure 3). Furthermore, we found no evidence for the presence of a functional duplicated *TLR5* copy in these genomes. Together, this shows that the observed losses of *TLR5* are not artifacts arising from aligning paralogous or processed pseudogenes.

Third, since no raw sequencing reads were available to confirm the four *TLR5*-inactivating mutations observed in the Yangtze river dolphin assembly (Figure 1A), we made use of the recently assembled genomes of the Amazon river and La Plata dolphins, which are sister species of the Yangtze river dolphin (Geisler, et al. 2011), and investigated whether any of these inactivating mutations are shared. Indeed, we found that the second stop codon mutation is shared between all three river dolphin species (Figure 1A). The presence of the same mutation in independently sequenced and assembled genomes makes it extremely unlikely that this mutation is a base error. Instead, this shared mutation strongly indicates that *TLR5* was already lost in the common ancestor of the three river dolphin species. We also found additional species-specific frameshifting mutations in the Amazon river and La Plata dolphins, which further support that *TLR5* is lost in these two species as well (Supplementary Figure 4). Similarly, while no sequencing reads are available to confirm mutations in the Hawaiian monk seal assembly, we found that six of seven observed mutations are shared with the Weddell seal (Figure 1A), providing strong support that *TLR5* is lost in the Hawaiian monk seal.

To investigate whether *TLR5* loss is specific to the domestic guinea pig or whether this gene is also lost in wild guinea pig species, we analyzed the *TLR5* locus in the genomes of two additional guinea pigs which are not a part of our genome alignment - Brazilian and Montane guinea pigs. As shown in Figure 1A, both stop codons and the 2 bp deletion are shared between all three guinea pig species. Additionally, the large deletion that removes most of the *TLR5* coding exon has the same breakpoints in all three guinea pig species (Figure 1B), which makes an assembly error highly unlikely and suggests that this deletion already occurred in their common ancestor. Furthermore, all three guinea pig species share larger structural rearrangements in this locus including several duplications that happened after the deletion (Supplementary Figure 5).

*TLR5* is entirely deleted in the genomes of the Chinese and Malayan pangolin (Figure 1A). To further investigate whether this deletion could be an assembly error, we generated alignment chains for the Tree pangolin, another recently sequenced pangolin species that was not included in our alignment. We found that *TLR5* is also deleted in the tree pangolin and the deletion break points are shared between all three pangolins (Figure 1C), which strongly suggests that the gene deletion already occurred in the common ancestor of the three pangolin species.

The three pinniped species in our data set share several frameshifting deletions (Figure 1A). Since these three species represent only the two of the three basal pinniped lineages, Phocidae (Weddell seal, Hawaiian monk seal) and Odobenidae (walrus), we investigated whether *TLR5* is also lost in the third pinniped lineage Otariidae. To this end, we used PCR to amplify *TLR5* from tissue samples of the brown fur seal, an otariid seal. Sequencing confirmed that the 25 bp and 17 bp deletion mutations are also present in the brown fur seal (Figure 1D). Furthermore, we found the same mutations in the Antarctic fur seal genome (Supplementary Figure 6). Together, the presence of inactivating mutations shared between Otariidae, Phocidae and Odobenidae suggests that *TLR5* loss already occurred in the pinniped stem lineage at least ~26 Mya (the estimated split of pinnipeds according to TimeTree (Kumar, et al. 2017b)).

Fourth, we used available transcriptomic data to explore whether there is still expression at the *TLR5* locus in the gene loss species. While we found no evidence for expression in the guinea pig and pangolin (Supplementary Figures 7, 8), clear expression at the *TLR5* locus was observed in Weddell seal (Supplementary Figure 9). Importantly, RNA-seq reads exhibit the frameshifting deletions that are present in the Weddell seal genome assembly, showing that these transcripts cannot encode a functional TLR5 protein. Consistent with previous findings (Sadier, et al. 2018), this suggests that the remnants of a lost coding gene can remain to be transcribed, in our case for at least 26 My.

Finally, we used RELAX (Wertheim, et al. 2015) to test whether *TLR5* evolves under relaxed selection in pinnipeds and the river dolphin where a substantial portion of the gene is still present. We did not apply RELAX to pangolins and guinea pigs where >80% of the gene is deleted. Further supporting *TLR5* loss in pinnipeds and Yangtze river dolphin, we found significant evidence for relaxed selection in both lineages (Ka/Ks = 1.17, P-value = 0.0002 for Yangtze river dolphin and Ka/Ks = 0.78, P-value = 0.0009 for pinnipeds) (Supplementary Table 3).

With several lines of evidence ranging from signatures of relaxed selection and the presence of validated inactivating mutations, we conclusively show that *TLR5* has been lost in four independent mammalian lineages. These mutations either delete the majority or the entire gene (guinea pigs, pangolins) or severely affect the reading frame and domains required for protein function (Figure 1A), suggesting that the remnants of *TLR5* cannot encode a functional flagellin-recognition receptor anymore.

Apart from TLR5, mouse Tlr11 has been shown to bind flagellin (Mathur, et al. 2012), even though the Tlr11-flagellin interaction is restricted to acidic conditions (Hatai, et al. 2016). Therefore, we considered the possibility that TLR5 loss may be compensated by the presence of TLR11, and investigated whether the *TLR5*-loss species possess an intact *TLR11* gene. However, we found that *TLR11* is most likely lost in the genomes of guinea pig, river dolphin, pinnipeds, and pangolins (Supplementary Figure 10), indicating that this gene cannot compensate for the loss of the flagellin-sensing *TLR5*.

## Discussion

Here, we show that *TLR5*, the gene encoding the major extracellular flagellin receptor, has been convergently lost four times in mammalian evolution. Our study highlights the importance of carefully validating putative gene-inactivating mutations to detect real gene loss events. All frameshifts detected by our gene loss detection method (Sharma, et al. 2018a) are really present in the respective assemblies, corroborating the high accuracy of our method; however, our sequencing read validation revealed that many of these frameshifts are actually base errors in genome assemblies that were generated from short or long sequencing reads. While current genome assembly efforts often focus on generating highly-contiguous assemblies (Miga, et al. 2019), our observations suggest that assembly base accuracy is potentially an underappreciated issue. Indeed, near-perfect assembly base accuracy would be of great importance to automatically analyze genomic data without the need to manually validate putative mutations.

Based on human and mouse studies (Hawn, et al. 2003; Feuillet, et al. 2006; Uematsu, et al. 2006; Vijay-Kumar, et al. 2007), loss of *TLR5* is predicted to severely impair the capacity to sense extracellular flagellin, in particular because all four *TLR5*-loss lineages also lack an intact *TLR11* gene that may have acted as a compensating factor. An intriguing question is why the evolutionary inactivation of both extracellular flagellin receptors does apparently not have deleterious consequences for guinea pigs, river dolphins, pinnipeds, and pangolins. A potential explanation could be that alternative pathways for flagellin detection compensate for the loss of *TLR5* and *TLR11*. For example, we recently described mammals that lost several human disease-associated genes but do not show the deleterious, disease-resembling phenotypes (Sharma, et al. 2019), which suggests the existence of either functionally-redundant genes or compensatory mechanisms that may have permitted losses of human disease genes in other mammals.

Whether the “natural *TLR5* knockout” mammals retain a capacity to sense extracellular flagellin could be tested experimentally. While research in river dolphins, pangolins and pinnipeds is generally impractical, guinea pigs have been used in biomedical research including studies of asthma, tuberculosis, Zika virus and other infectious diseases (Meurs, et al. 2006; Clark, et al. 2014; Kumar, et al. 2017a), because many aspects of guinea pig physiology and immunology are more similar to humans than to mouse or rat (Padilla-Carlin, et al. 2008). Thus, in addition to targeted research in mouse, guinea pigs could be used as another rodent model to study the consequences of natural *TLR5* knockout mammals.

Our findings raise questions of which evolutionary and ecological factors are associated or drove the evolutionary losses of *TLR5* in non-human mammals. *TLR5* also plays an important role in establishing and maintaining a healthy gut microbiome (Vijay-Kumar, et al. 2010; Fulde, et al. 2018) and gut microbiome composition is influenced by diet (Delsuc, et al. 2014; Youngblut, et al. 2019), raising the possibility that dietary switches could be one of the involved factors. However, while all four mammalian *TLR5*-loss lineages are dietary specialists, they specialize in different diets, ranging from carnivorous (river dolphins, pinnipeds), herbivorous (guinea pigs) to myrmecophagous (pangolins that feed on termites and ants) diets. This indicates that specialization to a particular diet was likely not a major factor behind the evolutionary losses of this gene. Furthermore, many other mammals convergently specialized on carnivorous, herbivorous and myrmecophagous diets without losing *TLR5*.

In general, loss of a gene can be due to relaxed selection to preserve its function or provide an evolutionary advantage (Albalat, et al. 2016; Sharma, et al. 2018a; Hecker, et al. 2019b; Huelsmann, et al. 2019). Interestingly, in humans, the *TLR5* stop codon mutation is associated with both positive and negative consequences. On the one hand, *TLR5* inactivation is associated with an increased susceptibility to Legionella-induced pneumonia and recurrent urinary tract infections, and constitutes a risk factor for type 2 diabetes (Hawn, et al. 2003; Hawn, et al. 2009; Al-Daghri, et al. 2013). On the other hand, *TLR5* inactivation is associated with an improved survival in melioidosis patients (West, et al. 2013), and in certain ethnic groups with a protection from systemic lupus erythematosus and Crohn’s disease (Hawn, et al. 2005; Gewirtz, et al. 2006). This indicates that inactivating *TLR5* can also have a protective effect against the immune system dysregulation that is involved in these diseases. Such beneficial effects may explain why this stop codon mutation has reached a relatively high frequency of >4% in many human populations, although signatures of recent adaptive evolution have not been detected (Barreiro, et al. 2009; Wlasiuk, et al. 2009). Since an inflammatory response represents a cost–benefit trade-off that is often optimized for a particular environment (Okin, et al. 2012), adjusting inflammatory responses can be an advantageous evolutionary strategy for species occupying specific environmental niches, exemplified by an attenuated inflammatory response to lipopolysaccharides in blood of deep diving seals (Bagchi, et al. 2018). Thus, it remains to be elucidated whether the *TLR5* losses in the four non-human mammalian lineages were due to relaxed selection or offered an evolutionary advantage.

## Materials and Methods

### Investigating and validating losses of *TLR5*

We used our gene loss detection method (Sharma, et al. 2018a) to investigate whether gene-inactivating mutations (frameshifting insertions/deletions, premature stop codon mutations, splice site disrupting mutations, large deletions) occur in *TLR5* in mammals. Since the two *TLR5* transcripts in human GENCODE version 32 (Harrow, et al. 2012) annotate an identical single coding exon and only differ in 5’ UTR exons, we focused on this single coding exon. We used human as the reference species and initially considered a total of 119 non-human mammals that are part of a 120-mammal genome alignment (Hecker, et al. 2020). The *TLR5* sequence provided by this alignment was re-aligned with CESAR (Codon Exon Structure Aware Realigner) (Sharma, et al. 2016; Sharma, et al. 2017) to avoid spurious frameshifts due to alignment ambiguities. Assembly gaps that overlap parts of *TLR5* in two species were not taken as evidence for gene loss (see Supplementary Figure 11).

Validation of small putative inactivating mutations was done as described before (Jebb, et al. 2018; Sharma, et al. 2018b). Briefly, we extracted a 50 bp genomic context surrounding a mutation, aligned these sequences against sequencing reads stored in SRA (Kodama, et al. 2012) (accession numbers for each species are listed in Supplementary Table 1), and counted how many reads support the inactivating mutation or the ancestral non-gene-inactivating allele. To investigate large deletions, we visualized and inspected pairwise alignment chains (Kent, et al. 2003) in the UCSC genome browser (Haeussler, et al. 2019). In addition, alignment chains were used to investigate whether remnants of *TLR5* occur in a context of conserved gene order and to exclude the possibility that a functional *TLR5* copy exists elsewhere in these assemblies.

Gene-inactivating mutations that are shared between related species are strong evidence against assembly base errors and indicate that gene loss is fixed in the respective clade (Hecker, et al. 2019b; Huelsmann, et al. 2019; Sharma, et al. 2019). Therefore, in addition to the 119 non-human mammals included in the 120-mammal genome alignment, we analyzed *TLR5* in six additional mammals that are close sister species to *TLR5*-loss species (two river dolphins, two guinea pigs, tree pangolin and Antarctic fur seal; NCBI assembly accession numbers are listed in Supplementary Table 1). For these six species, we computed sensitive pairwise alignment chains to human (hg38) using lastz (Harris 2007) (alignment parameters K = 2400, L = 3000, Y = 9400, H = 2000, default scoring matrix), axtChain (Kent, et al. 2003) (linearGap=loose, otherwise default parameters), and RepeatFiller (Osipova, et al. 2019) (default parameters). The presence of shared inactivating mutations was manually confirmed.

### Relaxed selection

We used RELAX (Wertheim, et al. 2015) to determine whether the remaining *TLR5* coding sequence evolves under relaxed selection in the Yangtze river dolphin and in pinnipeds. To this end, we first extracted the coding sequence of *TLR5* from the 120-mammal alignment. Then, we generated pairwise alignments between the human coding sequence and the sequence corresponding to each of the 119 mammals using CESAR. These pairwise alignments were combined into a multiple sequence alignment using maf-join (Kielbasa, et al. 2011). The multiple alignment was further post-processed by removing alignment columns that contained frameshifting insertions and by substituting in-frame stop codons by NNN. We ran the test for relaxed selection twice and specified either the Yangtze river dolphin branch or the branch leading to pinnipeds as the foreground. All other branches leading to species that did not lose *TLR5* were specified as background.

### Analysis of transcriptomic data

RNA-seq data of guinea pig brain (SRP017611) (Fushan, et al. 2015), lung (SRP040447) (Davidsen, et al. 2014), ovaries and testis (SRP104222) (Bens, et al. 2018), Malayan pangolin cerebrum and lung (SRP064341) (Mohamed Yusoff, et al. 2016), small intestine, large intestine and stomach (SRP101596 and SRP156258, Guangdong Institute of Applied Biological Resources), and Weddell seal brain, lung, placenta, and testis (SRP200409, Broad Institute) was downloaded from the Sequence Read Archive (SRA) (Kodama, et al. 2012). Next, fastq-dump was used to process the initial read data using parameters for removing technical reads (skip-technical), filtering low-quality reads (read-filter pass), removing tags (clip), converting data into base space (dumpbase), keeping read identifiers (readids) and splitting paired-end reads into separate files (split-files). Reads were mapped to the corresponding genomes with STAR (version 2.4.2a) (Dobin, et al. 2013). For Malayan pangolin, we adjusted the number of bins for genomic indexes (genomeChrBinNbits: 15). Reads were mapped separately per run as paired-end reads with parameters limiting the ratio of mismatches per mapped read (outFilterMismatchNoverLmax=0.04) and limiting the number of mapping locations per read (outFilterMultimapNmax=20), and otherwise defaults. To visualize gene expression in the UCSC Genome Browser (Haeussler, et al. 2019), we processed the mapped reads with BEDtools (Quinlan, et al. 2010) and bedGraphToBigWig (Kent, et al. 2010).

### PCR analysis of TLR5 in the brown fur seal

Tissue samples of the brown fur seal (*Arctocephalus pusillus; Pinnipedia, Otariidae*) were obtained from Senckenberg Natural History Collections Dresden, Germany (collection ID: MTD-TB 250) representing a specimen from the Zoological Garden Leipzig (Germany). DNA was extracted using the innuPrep DNA Mini Kit (Analytik Jena, Jena, Germany) following the manufacturer’s instructions (protocol for DNA isolation from tissue samples or rodent tails) except that tissue lysis was performed overnight and DNA was precipitated in two steps using 50μl elution buffer. Standard PCR reactions were performed in a total volume of 20μl using 10-20ng DNA, one unit DFS-Taq polymerase (Bioron, Ludwigshafen, Germany) in the recommended buffer, 3.5 mM MgCl_2_, 0.2 mM of each dNTP (Fermentas, St. Leon-Rot, Germany), and 0.37 μM of each primer listed in Supplementary Table 4. PCR amplification of target *TLR5* regions was performed with primer combinations shown in Supplementary Figure 12, using 35 PCR cycles with denaturation at 94°C (15 sec but 3 min for the first cycle), annealing at 63°C (20 – 30 sec), and extension at 72°C (1:30 min but 7 min for the last cycle). PCR products were Sanger sequenced in both directions after enzymatic clean-up with ExoSAP-IT (USB Europe GmbH, Staufen, Germany), cycle sequencing using the PCR primers, the BigDye Terminator v3.1 Cycle Sequencing Kit (Applied Biosystems) and an ABI 3130xl Genetic Analyser (Applied Biosystems, Foster City, CA, USA).

## Data availability

All analyzed genome assemblies and sequencing read data is publicly available on NCBI and SRA (accession numbers listed in Supplementary Table 1). The multiple genome alignment is available at https://bds.mpi-cbg.de/hillerlab/120MammalAlignment/Human120way/. The *TLR5* sequence of the brown fur seal has been submitted to NCBI (accession number pending).

## Acknowledgment

We thank the genomics community for sequencing and assembling the genomes and the UCSC genome browser group for providing software and genome annotations. We also thank Bogdan Kirilenko for computing river dolphin genome alignments, and the Computer Service Facilities of the MPI-CBG and MPI-PKS for their support. This work was supported by the Max Planck Society, the German Research Foundation (HI 1423/3-1) and the Leibniz Association (SAW-2016-SGN-2).

