## Supplementary Material for "Convergent losses of *TLR5* suggest altered extracellular flagellin detection in four mammalian lineages"

This PDF file contains:

- Figures 1-12
- Supplementary References

Supplementary Tables 1-4 are provided as sheets in a separate Excel file.

A

```
                   Human     AACCCATTCAGAA
     Orangutan (ponAbe)      AACCCA-TCAGAA
  SRR748041.97763734.1       AACCCATTCAGAA
  SRR748041.74482427.2       AACCCATTCAGAA
  SRR748041.26867665.1       AACCCATTCAGAA
  SRR748042.38070946.2       AACCCATTCAGAA
  SRR748041.28432315.1       AACCCATTCAGAA
```

B

```
                   Human  TGGCTGGTTTCTT
        Alpaca (vicPac2)  TGGCTG-TTTCTT
  SRR1552599.200202768.1  TGGCTGGTTTCTT
  SRR1552599.184451509.2  TGGCTGGTTTCTT
  SRR1552599.158573862.1  TGGCTGGTTTCTT
  SRR1552599.149519331.1  TGGCTGGTTTCTT
  SRR1552599.139788638.1  TGGCTGGTTTCTT
```

C

```
                          Human  AGGACA-GTCACT
  Domestic sheep (HLoviAri4)     AGGACA-GTCAGA
   Bighorn sheep (HLoviCan1)     AGGACACGTCAGA
       SRR501895.162761708.2     AGGACA-GTCAGA
       SRR501895.149809378.1     AGGACA-GTCAGA
       SRR501895.149159580.1     AGGACA-GTCAGA
       SRR501895.119549498.2     AGGACA-GTCAGA
       SRR501895.119549498.1     AGGACA-GTCAGA
```

D

```
                          Human  AGGACA-GTCACT
  Domestic sheep (HLoviAri4)     AGGACA-GTCAGA
   Bighorn sheep (HLoviCan1)     AGGACACGTCAGA
        SRR501895.96850434.1     AGGACA-GTCAGA
        SRR501895.92310229.2     AGGACA-GTCAGA
        SRR501895.81373516.1     AGGACA-GTCAGA
        SRR501895.29601966.1     AGGACA-GTCAGA
        SRR501895.17753062.2     AGGACA-GTCAGA
```

E

```
                       Human  TTTAAA-GGCAAAT
  Little brown bat (myoLuc2)  CTTGCCCGTCTTTA
     SRR6793294.47404072.2    CTTGCC-GTCTTTA
     SRR6793294.45915763.1    CTTGCC-GTCTTTA
     SRR6793294.44209865.2    CTTGCC-GTCTTTA
     SRR6793294.43265995.1    CTTGCC-GTCTTTA
     SRR6793294.42772856.1    CTTGCC-GTCTTTA
```

F

```
                       Human  AAAGTCCATAGAT
  Little brown bat (myoLuc2)  GAAGTC-ATCGAC
     SRR6793295.51222580.2    GAAGTCCATCGAC
     SRR6793295.51177232.2    GAAGTCCATCGAC
     SRR6793295.51177232.1    GAAGTCCATCGAC
     SRR6793295.50504314.2    GAAGTCCATCGAC
     SRR6793295.50452996.1    GAAGTCCATCGAC
```

G

```
                  Human  AGAAAGATCGTTT
     Elephant (loxAfr3)  CGAAAG-TTGTTT
  SRR958468.52800368.2   CGAAAGGTTGTTT
  SRR958468.25619718.2   CGAAAGGTTGTTT
  SRR958468.49633226.2   CGAAAGGTTGTTT
  SRR958468.37749047.1   CGAAAGGTTGTTT
  SRR958468.38174150.2   CGAAAGGTTGTTT
```

H

```
                  Human  CCATCT-CCTAAT
     Elephant (loxAfr3)  CCATCTTCCTAGT
  SRR958467.88434749.2   CCATCT-CCTAAT
  SRR958467.84102042.1   CCATCT-CCTAAT
  SRR958467.64715860.2   CCATCT-CCTAAT
  SRR958467.42899696.2   CCATCT-CCTAAT
```

I

```
                   Human  TGAGGTGGCCTGAGG
     Armadillo (dasNov3)  TGAGGTG-CCGGAAG
  SRR6206918.67775986.1   TGAGGTGGCCGGAAG
  SRR6206918.67351711.2   TGAGGTGGCCGGAAG
  SRR6206918.64951246.1   TGAGGTGGCCGGAAG
  SRR6206918.19294686.1   TGAGGTGGCCGGAAG
  SRR6206918.18706253.2   TGAGGTGGCCGGAAG
```

J

```
                               Human  CTAGGAGTGGTGCTCATGGCCGGTC-CTGTGTTTGGAATTCCTTCC
                    Rabbit (oryCun2)  GCAGCAG-GGAGCTCGTGGCCAGCTGCCGGGTCTGGA-TTCTTACT
           RefSeq TLR5 (NM_001329083.1)  GTAGGAGTGGTGCTCCTGGCCAGCC-CCGGGTTTGGAATTTCTTCC
  A00489:19:H3GG3DRXX:1:2264:1470:18912  GTAGGAGTGGTGCTCCTGGCCAGCC-CCGGGTTTGGAATTTCTTCC
  A00489:19:H3GG3DRXX:1:2262:28302:3787  GTAGGAGTGGTGCTCCTGGCCAGCC-CCGGGTTTGGAATTTCTTCC
   A00489:19:H3GG3DRXX:1:2227:3269:6527  GTAGGAGTGGTGCTCCTGGCCAGCC-CCGGGTTTGGAATTTCTTCC
   A00489:19:H3GG3DRXX:1:2227:3242:6480  GTAGGAGTGGTGCTCCTGGCCAGCC-CCGGGTTTGGAATTTCTTCC
  A00489:19:H3GG3DRXX:1:2145:6804:18443  GTAGGAGTGGTGCTCCTGGCCAGCC-CCGGGTTTGGAATTTCTTCC
```

**Supplementary Figure 1:** Several putative inactivating mutations in *TLR5* are base errors in the genome assemblies.

Each panel show the human genomic sequence (hg38 assembly) around a putative mutation (red font) detected in the assembly of another mammal.

(A) 1 bp frameshifting deletion in orangutan.

(B) 1 bp frameshifting deletion in alpaca.

(C, D) 1 bp frameshifting insertions in bighorn sheep.

(E, F) 1 bp frameshifting insertion and deletion in little brown bat. We used RNA-seq reads to show that both frameshifts are base errors in the assembly. This is further supported by searching Sanger sequencing reads, which revealed that a single read (ti:988238460) with low quality scores around both frameshifts was likely incorporated into the genome assembly.

(G, H) 1 bp frameshifting insertion and deletion in elephant.

(I) 1 bp frameshifting deletion in armadillo. Since genomic read data is not available for the armadillo, we analyzed RNA-seq read data. Further corroborating that this frameshift is a base error in the assembly, we searched the Sanger sequencing reads stored in the NCBI trace archive, which revealed that the single read that was likely incorporated into the assembly (ti:561639686) has low quality scores around the region of the base error.

(J) Frameshifting deletions and insertions in rabbit. Searching the Sanger sequencing reads stored in the NCBI trace archive revealed that a single read (ti:1946048333) with extremely low quality scores around this region was likely incorporated into the rabbit genome assembly. Further demonstrating that these frameshifts are base errors, RefSeq annotates a *TLR5* gene with an intact reading frame (NM_001329083.1, based on a sequenced mRNA HQ874605.1), which is supported by RNA-seq read data.

A

```
                      Human AAGGACCATCCCCAGGGCA
Gorilla (gorGor5 assembly) AAGGACCAT-CCCAGGGCA
Gorilla (gorGor3 assembly) AAGGACCATCCCCAGGGCA
      SRR957682.59993119.2 AAGGACCATCCCCAGGGCA
      SRR957682.34059603.2 AAGGACCATCCCCAGGGCA
      SRR957682.10163173.2 AAGGACCATCCCCAGGGCA
      SRR957683.81806585.2 AAGGACCATCCCCAGGGCA
      SRR957683.21643372.2 AAGGACCATCCCCAGGGCA
```

B

```
                      Human GTTCCATCTGTTTGAACTT
   Pig (susScr11 assembly) GCCCCATCT-TCTGAACTT
    Pig (susScr3 assembly) GCCCCATCTTTCTGAACTT
      SRR652354.24987092.2 GCCCCATCTTTCTGAACTT
      SRR652353.50513659.2 GCCCCATCTTTCTGAACTT
      SRR652352.32451792.1 GCCCCATCTTTCTGAACTT
      SRR652352.14449607.2 GCCCCATCTTTCTGAACTT
      SRR652352.11059424.1 GCCCCATCTTTCTGAACTT
```

**Supplementary Figure 2:** Erroneous frameshifting deletions in the *TLR5* gene that were introduced in the latest PacBio-based gorilla and pig genome assemblies.

(A) The gorilla gorGor5 assembly (Gordon, et al. 2016) shows a 1 bp frameshifting deletion (red font, highlighted by the arrow). However, this frameshift is an error, likely introduced by PacBio reads that were used to assemble gorGor5, as neither the previous short read-based gorGor3 assembly nor raw Illumina reads support this frameshift. For space reasons, only 5 reads are shown.

(B) Similar to (A), the recent PacBio-based susScr11 pig assembly (Warr, et al. 2019) shows a 1 bp frameshift in a different position in *TLR5*. Since neither the previous susScr3 assembly nor raw Illumina reads support this base deletion, this frameshift is an error newly introduced in susScr11.

These examples support that insertion/deletion errors can be a frequent type of base errors in long read-based assemblies (Watson, et al. 2019).

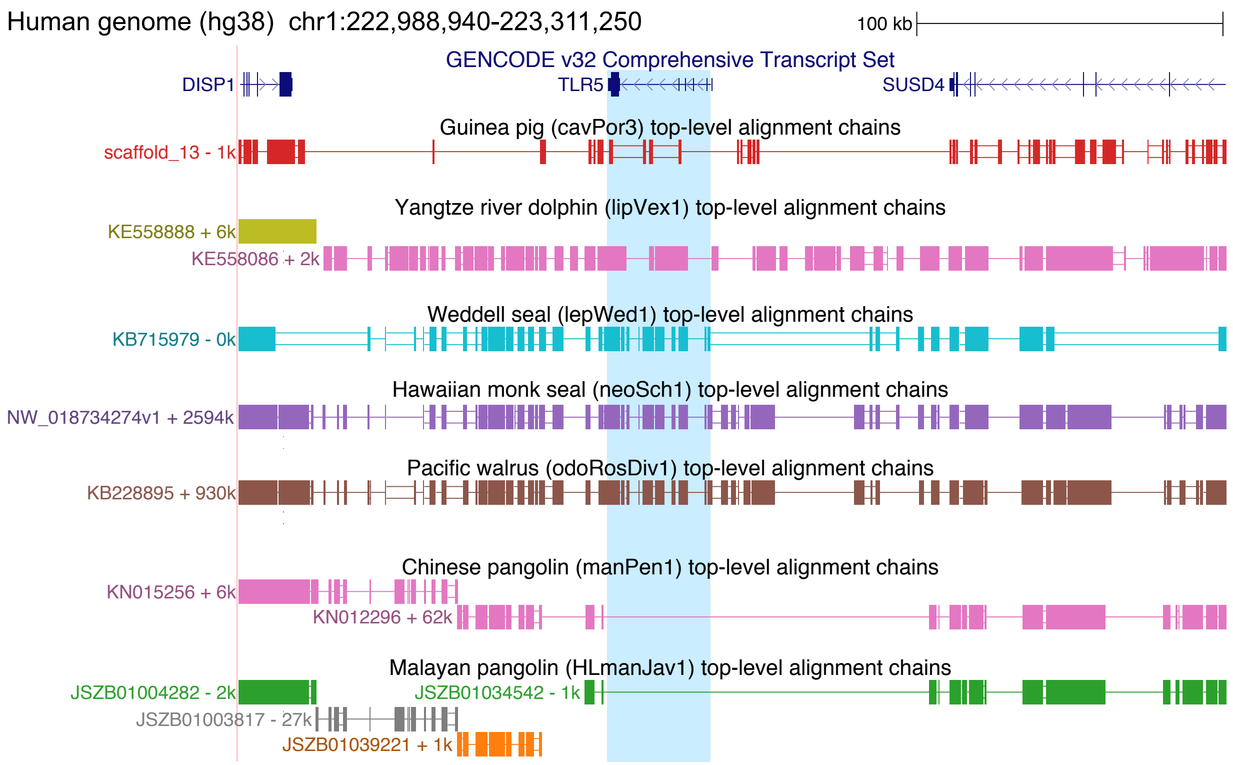


**Supplementary Figure 3:** Remnants of *TLR5* occur in the context of conserved gene order.

UCSC genome browser screenshot shows the human *TLR5* locus and the top-level chain of co-linear local alignments to species that lost the *TLR5* gene (highlighted in light blue). Blocks in an alignment chain represent aligning regions, double lines represent unaligning sequence, and single lines represent deletions. For all species, the alignment chain that spans *TLR5* also aligns upstream intergenic sequence and the downstream *SUSD4* gene, showing that the remnants of *TLR5* occur in a context of conserved gene order. Furthermore, for the more contiguous assemblies of the guinea pig and the three pinnipeds, the alignment chain also aligns the upstream *DISP1* gene. Since the orthologous 322 kb locus shown in this figure is not assembled as a single but several scaffolds in the more fragmented river dolphin and pangolin genomes and since by definition a single chain cannot span different scaffolds, alignments upstream of *TLR5* are represented as separate chains for the river dolphin and pangolin genomes.

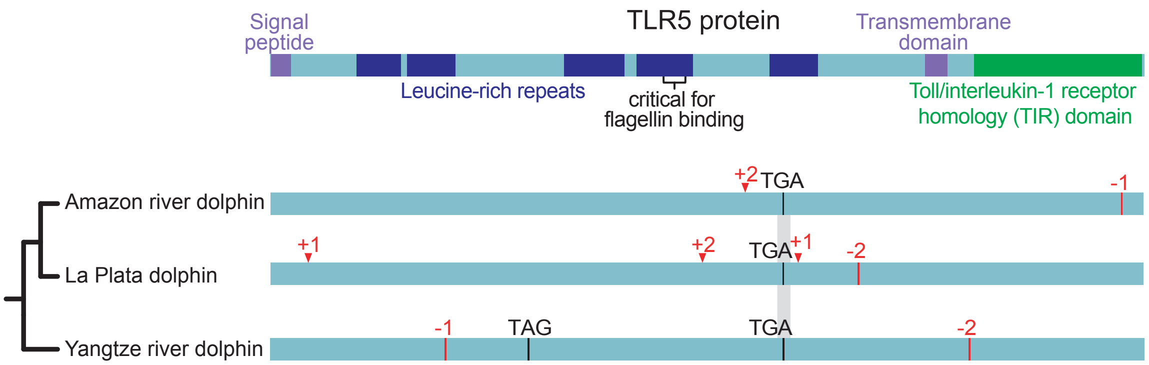


**Supplementary Figure 4:** Loss of *TLR5* in additional river dolphin species. Visualization of gene-inactivating mutations in the single coding exon of *TLR5* in the Amazon river and La Plata dolphin. The visualization of *TLR5* of the Yangtze river dolphin is shown again for comparison. The phylogeny of the three species is according to (Geisler, et al. 2011). While a stop codon mutation is shared between the three river dolphin species (grey background, see Figure 1A), Amazon river and La Plata dolphins show additional species-specific mutations, supporting that *TLR5* is also lost in these two species.

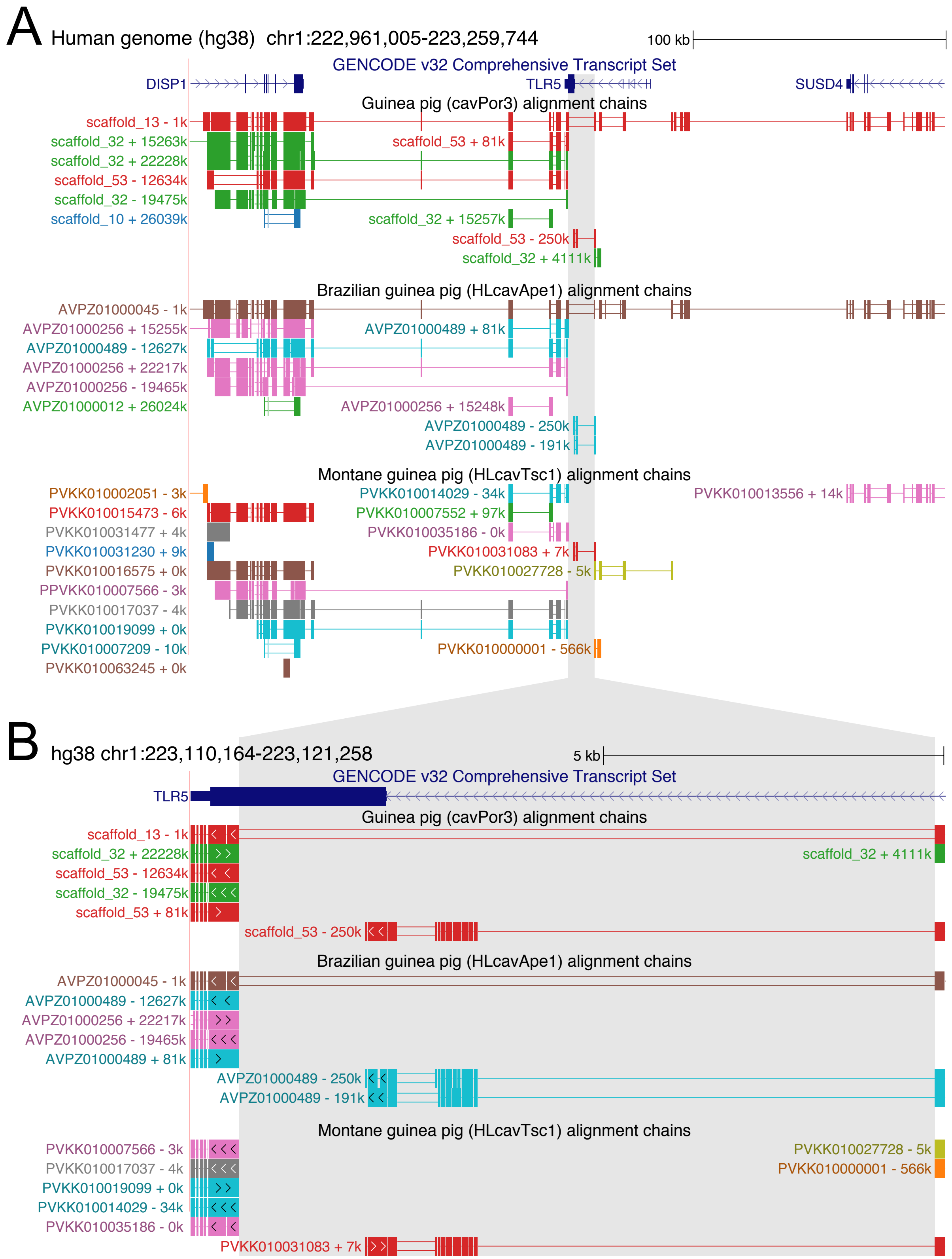


**Supplementary Figure 5:** The deletion of a large part of *TLR5* is shared among three guinea pig species.

UCSC genome browser screenshot shows the human *TLR5* locus with both flanking genes and alignment chains to three guinea pig species (A). The grey highlighted region highlights the large deletion that removes most of the coding exon of *TLR5*. The magnified view in (B) shows that this deletion has the same breakpoints in the domestic and Brazilian guinea pig, and very likely also in the more fragmented Montane guinea genome. In addition to this deletion, alignment chains suggest that several duplications comprising a part of the *DISP1* gene and the part of *TLR5* occurred before the split of these three guinea pigs. Since the downstream end of the duplicated region is similar to the deletion breakpoint, it is likely that these duplications that happened after the deletion.

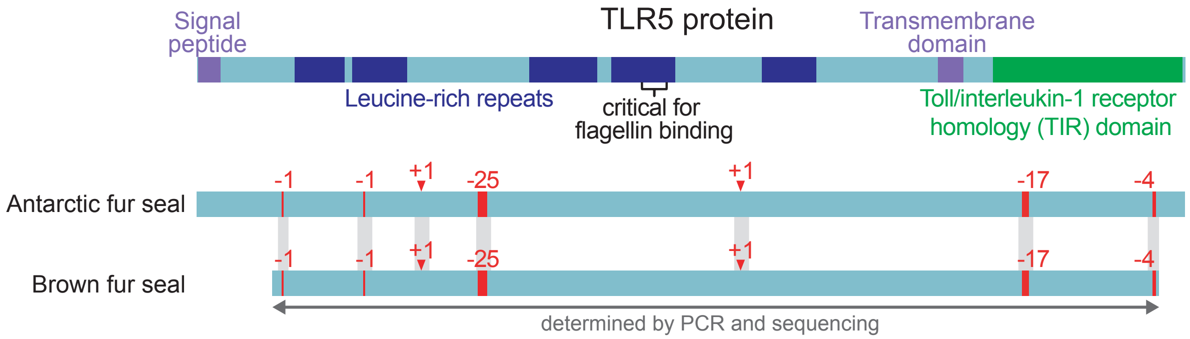


**Supplementary Figure 6:** Loss of *TLR5* in the Antarctic fur seal.
After we used PCR to sequence *TLR5* in the brown fur seal (shown again for comparison), a genome assembly of the Antarctic fur seal became available. Analyzing *TLR5* in the Antarctic fur seal, we found that the 1, 25 and 17 bp frameshifting deletions are shared between both fur seals (grey background). In addition, the Antarctic fur seal exhibits several lineage-specific mutations.

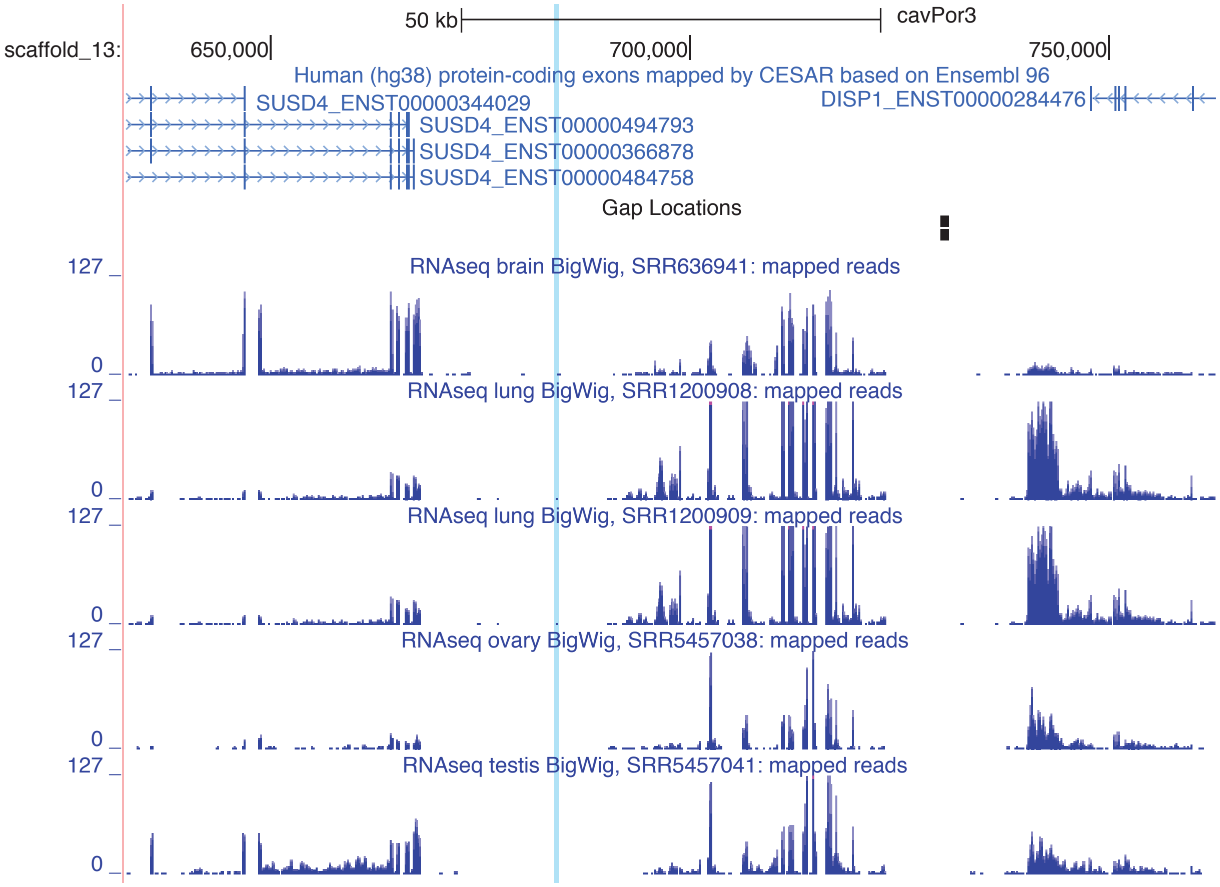


**Supplementary Figure 7:** Analysis of gene expression at the *TLR5* locus of guinea pig. For tissues that typically show clear expression of *TLR5* in human, mouse or rat, we mapped and analyzed available guinea pig RNA-seq data. The remnant of *TLR5* was inferred from the genome alignment and is highlighted in light blue. Assembly gaps are indicated by black blocks. In contrast to adjacent genes, *TLR5* exhibits no expression in any of the analyzed tissues in the guinea pig.

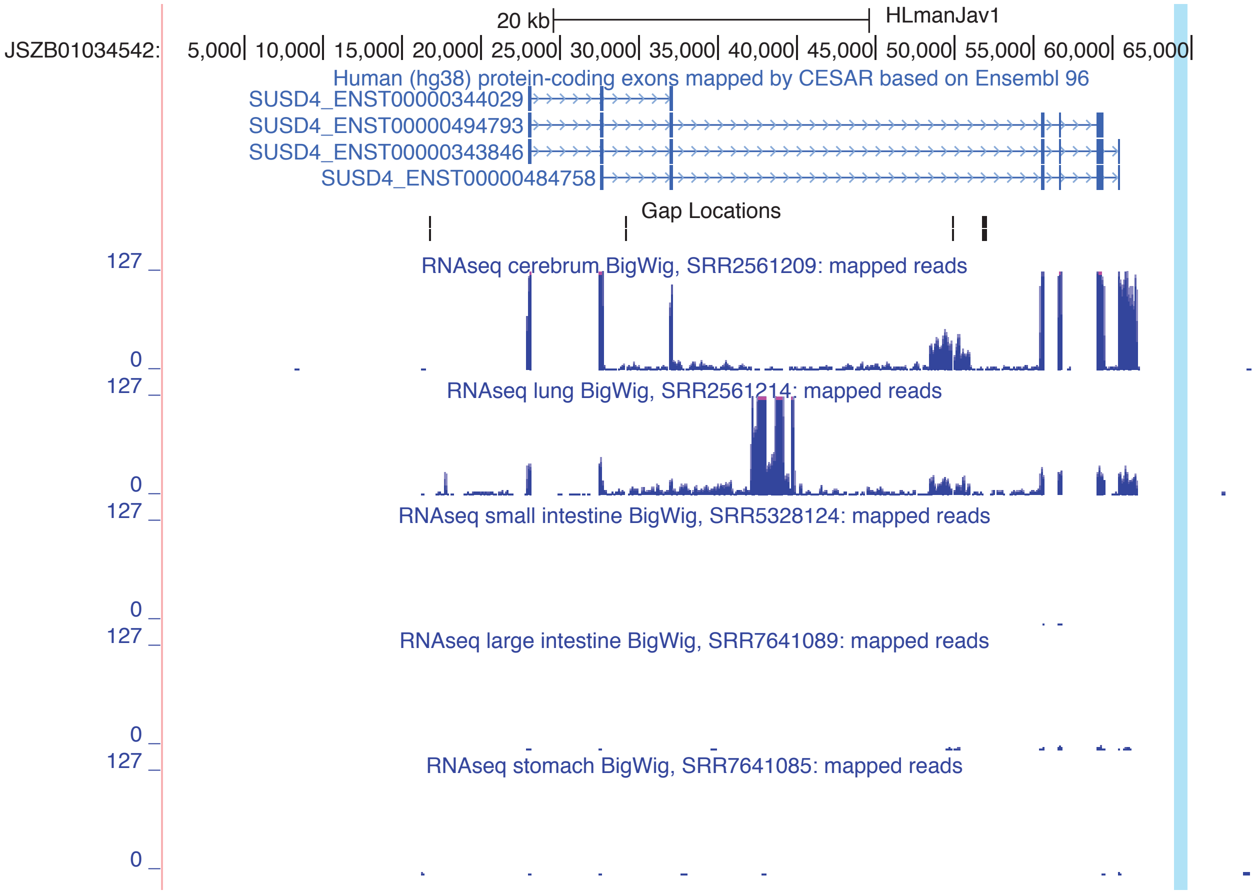


**Supplementary Figure 8:** Analysis of gene expression at the *TLR5* locus of Malayan pangolin.

For tissues that typically show clear expression of *TLR5* in human, mouse or rat, we mapped and analyzed available RNA-seq data of the Malayan pangolin. *TLR5* is completely deleted in pangolins and the locus around the deleted *TLR5* gene was inferred from the breakpoints (aligning sequence next to the deletion) based on the genome alignment and is highlighted in light blue. The entire scaffold that contains the *TLR5* locus in the Malayan pangolin assembly is shown. Assembly gaps are indicated by black blocks. In contrast to the adjacent *SUSD4* gene, *TLR5* is not expressed in any of the analyzed tissues in the Malayan pangolin.

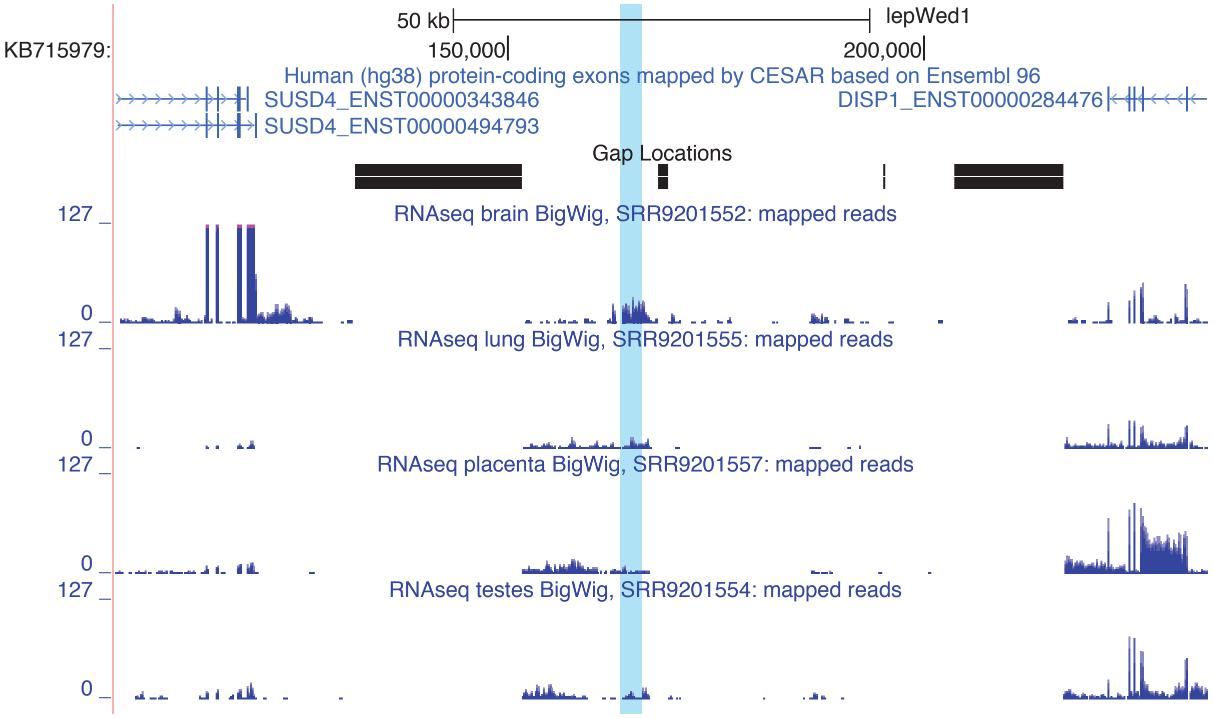


**Supplementary Figure 9:** Analysis of gene expression at the *TLR5* locus of Weddell seal.

For tissues that typically show clear expression of *TLR5* in human, mouse or rat, we mapped and analyzed available RNA-seq data of the Weddell seal. The remnant of *TLR5* was inferred from the genome alignment and is highlighted in light blue. Assembly gaps are shown as black blocks. We found that *TLR5* is expressed in brain of Weddell seal and lowly expressed in lung, placenta and testes. Importantly, while RNA-seq reads indicate that the most downstream -1 bp frameshifting deletion (see Figure 1A) may be polymorphic, other frameshifting deletions that are present in the genome are also present in the RNA-seq reads that align to the *TLR5* locus. This indicates that the respective transcripts cannot encode a functional toll-like receptor protein anymore.

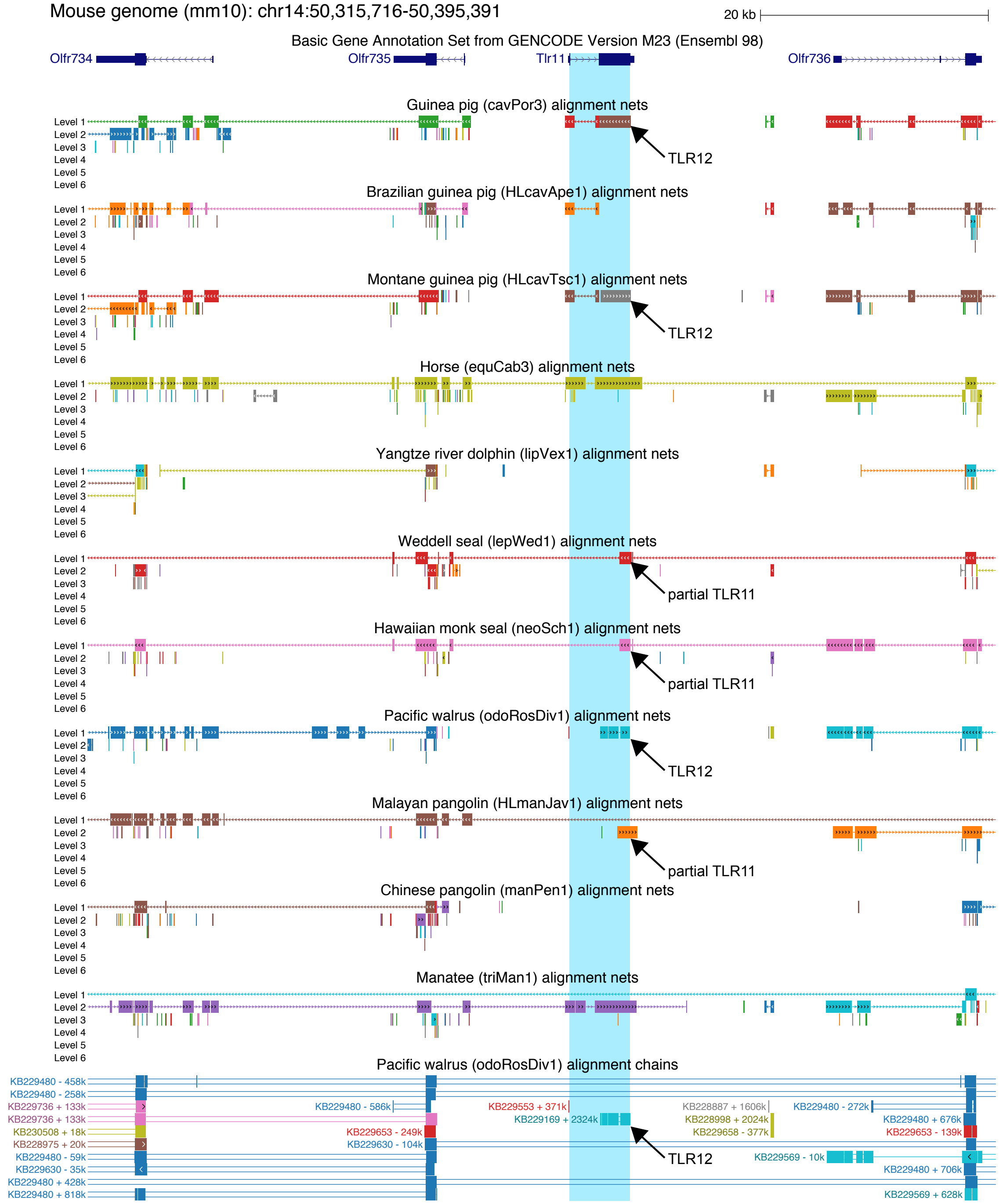


**Supplementary Figure 10:** *TLR5*-loss species do not possess an intact *Tlr11* gene. UCSC genome browser screenshot shows the mouse *Tlr11* locus and the alignment nets of *TLR5*-loss species, in addition to horse and manatee. *Tlr11* is located inside an olfactory receptor gene cluster. Since olfactory receptor genes have hundreds of paralogs and alignment chains capture both orthologs and paralogs (illustrated for a few walrus chains at the bottom), it is not possible to visualize alignment chains for all species. Therefore, we show alignment nets instead that typically represent orthologous alignments (Kent, et al. 2003).

For Brazilian guinea pig, Yangtze river dolphin and Chinese pangolin, the nets show that no genomic locus aligns to the large coding exon of *Tlr11*, indicating that the gene is not present in these species. For the domestic and Montane guinea pig, and Pacific walrus the best alignment is the paralog *Tlr12* (indicated by labeled arrows), which also indicates complete absence of *Tlr11*. For the Weddell seal, Hawaiian monk seal and Malayan pangolin, only a fragment of *Tlr11* was detected (indicated). In contrast, horse and manatee are two species where the entire *Tlr11* and the surrounding olfactory receptor genes align, showing that *Tlr11* originated before the split of placental mammals.

To confirm that none of the *TLR5*-loss species has a full length, intact *Tlr11* gene, we used the protein sequence of the Horse Tlr11 and performed sensitive tblastn searches (word-size = 2, gap-existence = 11, gap-extension = 1) against the genome assemblies of these species at NCBI. None of these searches returned a full-length Tlr11. Consistent with the genome alignments, we obtained a protein annotated as Tlr12 for guinea pigs and the partial *Tlr11* for Weddell seal, Hawaiian monk seal, and pangolins. These observations strongly indicate that *Tlr11* is lost in all *TLR5*-loss species.

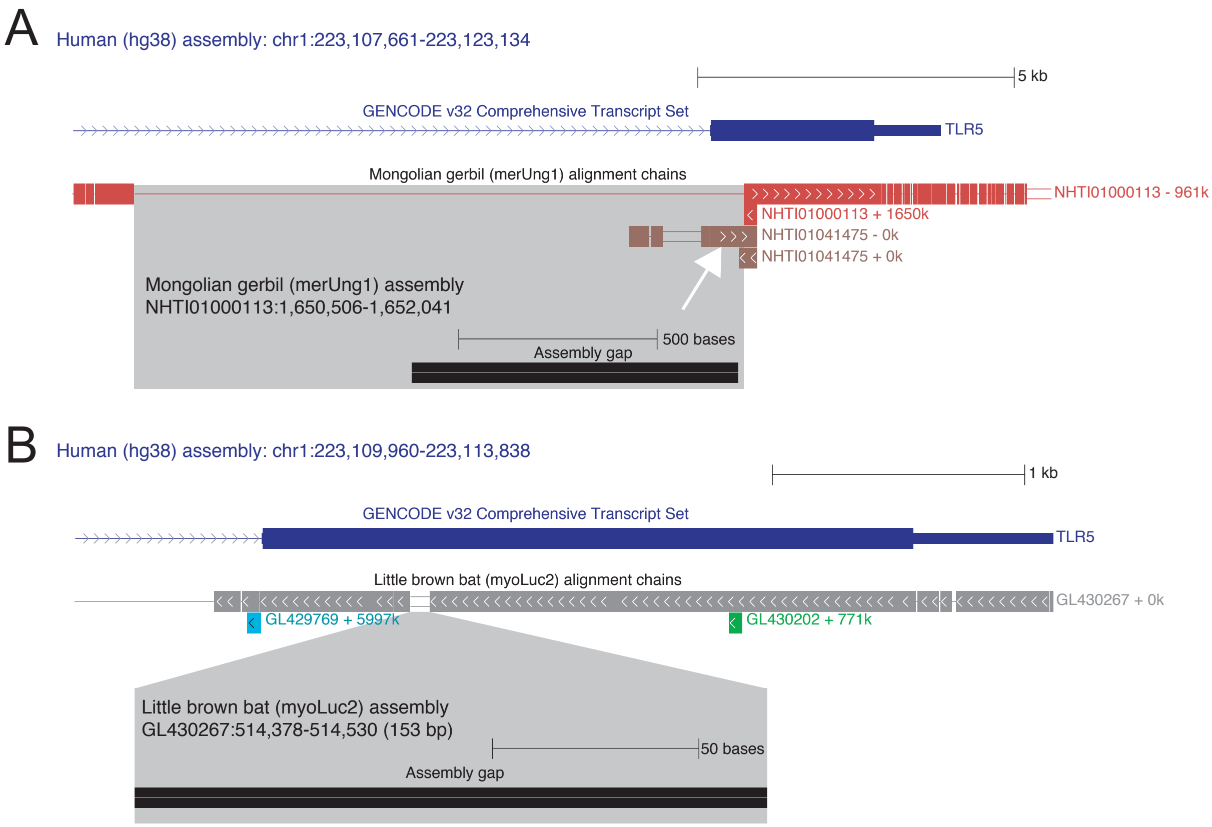


**Supplementary Figure 11**: Assembly gaps overlapping the *TLR5* loci in the Mongolian gerbil and little brown bat.

(A) In the Mongolian gerbil, the alignment of the single coding exon of *TLR5* are split and align to two genomic loci, represented by two separate alignment chains. The majority of the *TRL5* coding exon aligns to a 1.6 Mb long scaffold NHTI01000113 (red chain). However, the gerbil locus between the aligning blocks highlighted in grey overlaps an assembly gap that covers the 5' part of *TRL5*. This 5' *TLR5* part aligns to a separate contig NHTI01041475 (brown chain highlighted by a white arrow) that is just 1,972 bp long. Such assembly gaps should not be considered as evidence for a partial gene deletion and gene loss.

(B) A small region of *TLR5* does not align to the little brown bat; however, the respective locus entirely overlaps an assembly gap, which is not evidence for gene loss. Consistent with this, blasting the sequence of this assembly-gap-overlapping locus of related *Myotis* bats against RNA-seq data (SRX3752326) reveals several reads with 95% identity that span this assembly gap.

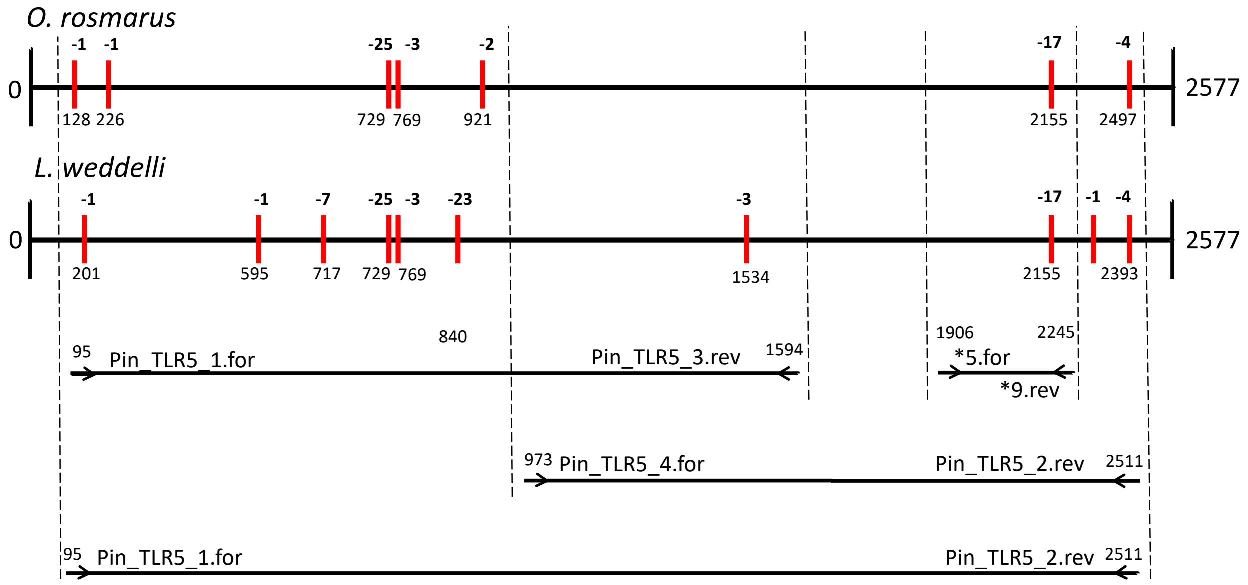


**Supplementary Figure 12:** Primer locations to sequence *TLR5* in the Brown fur seal. The figure illustrates the single *TLR5* coding exons of the Pacific walrus and Weddell seal, which we used to design primers to sequence *TLR5* in the Brown fur seal. Primer sequences are listed in Supplementary Table 4.
